# Optimising 3D scaffold for otic neural progenitor differentiation

**DOI:** 10.1101/120279

**Authors:** Kamolchanok Ngamkham, Marcelo N. Rivolta, Giuseppe Battaglia

## Abstract

Hearing loss is a condition highly prevalent worldwide. It affects people of a broad age range since the causes and risk factors are varied. At present, some types of hearing impairments have a palliative treatment whereas some, especially for those where otic neurons are damaged, cannot be properly treated. Recent findings had shown it possible to use human embryonic stem cell-derived otic neural progenitors (ONPs) as a new mode of treating hearing loss caused by damage to the spiral ganglion neurons (SGNs). To improve the efficiency and overcome some limitations of this potential treatment, we have applied principles of tissue engineering which involves an interaction between cells and an extracellular matrix –mimicking scaffold. Here, we describe the influence of poly(l-lactic acid)(PLLA) aligned fibres on ONP cell morphology, proliferation, neuronal differentiation and establishment of neural polarity under both progenitor and neuralising conditions. The results show that most of ONPs on aligned fibres exhibited bipolar morphology and extended their neurites along the major fibre axis. Their proliferation was lower than those in 2D culture but the differentiation of ONPs on aligned fibres was significantly enhanced in both progenitor and neuralising conditions as indicated by the fluorescence intensity and number of cells that were positive for neuronal markers (β-tubulin III and NF200) and the expression pattern of spiral ganglion molecular markers (MMP13, NPR2 and NTNG1). Moreover, axonal and dendritic markers (TAU and MAP2 respectively) were also induced after 14 days in culture.

## INTRODUCTION

Hearing loss is one of the most common causes of disability. According to the available data from the World Health Organization (WHO), around 360 million people worldwide are suffering from moderate or greater hearing loss. Several types of hearing loss can be categorised according to the area of the ear where the problem lies. Sensorineural hearing loss, which is caused by the damage to the hair cells or the auditory nerve in the inner ear is the most common. Unlike other nonmammalian vertebrates, damage or degeneration of human hair cells and spiral ganglion neurons (SGNs) is irreversible ^1,2^. Even though cochlear implants can be used to overcome a hair cell deficit, loss of SGNs can limit the effectiveness of this treatment ^3^. Thus, alternative strategies for SGN regeneration have been widely researched and stem cell therapy has been proposed as a potential therapeutic path.

Findings from a recent study highlight the possibility of SGNs restoration using a stem cell-based strategy. Otic neural progenitors (ONPs) that were differentiated from human embryonic stem cells were found to exhibit true otic characteristics as indicated by the expression of several otic markers such as PAX8, PAX2, SOX2 and FOXG1. They were also able to further differentiate into SGNs under ‘neuralising’ condition, becoming positive for neuronal markers, displaying appropriated membrane currents and firing action potentials. When transplanted into gerbils that were deafened by the application of ouabain (a model of auditory neuropathy), they engrafted and significantly improved the auditory thresholds ^4^.

However, there are some potential limitations in this study that might constrain a wider application of this approach. The deafness-inducing drug ouabain damages primarily the type I SGNs, while type II SGNs and glial cells remain mostly intact ^5^. These preserved tracks might act as a guidance cue for the transplanted cells, facilitating them to reach their targets centrally and on the periphery. In cases were these tracts are absent, such as in a cochlear nerve aplasia, ONPs might not be able to extend their neurites effectively and functional restoration might be impaired. Moreover, there are some putative drawbacks in delivering cells directly by cell injection. Based on evidence from other experimental systems, the retention of cells at the target site and cell survival rate can be poor ^6,7^. To overcome those limitations, the use of a support or tissue engineered scaffold mimicking the extracellular matrix may be advantageous.

The complexity of the cochlea and sound sensing mechanism make tissue engineering for the inner ear a difficult task ^8^. Most of the promising previous work relies on the improvement of the cochlear implant biological interface. Various reports indicated that alteration of electrode surface topography, chemistry or biological functionalisation offer a great opportunity for the improvement of this prostheses ^9–20^.

The physiological extracellular matrix of the inner ear has also been studied as a natural, tissue engineered scaffold. Mice, rat and human cochlea were decellularised and extracellular matrix contents, identified. However, these decellularised matrices will not be appropriated for clinical application of transplanted cells, and have not been widely used to support stem cell differentiation ^21^. In this study, we aimed to investigate the behaviour of ONPs on a biomaterial and explore if this scaffold could provide advantages for ONP transplantation. Poly(l-lactic acid)(PLLA), a biodegradable and biocompatible polymer, was fabricated as aligned electrospun fibres and used as a scaffold for ONP culture. To address the effect of aligned fibres on ONPs, cell attachment, proliferation, differentiation and establishment of neural polarity were then analysed.

## MATERIALS AND METHODS

### Human embryonic stem cell (hESC) culture

Shef1 hESC line was cultured and used for the derivation of otic neural progenitor. hESC colonies were maintained on inactivated mouse embryonic fibroblast (MEF) feeder layers on gelatin-coated tissue culture flasks. The culture media consisted of knockout Dulbecco’s modified Eagle’s medium (Invitrogen), 20% knockout serum replacement (KOSR), 1% nonessential amino acids, 2mM L-glutamine, (all from Invitrogen), 0.1mM ß-mercaptoethanol (Sigma), and 4ng/ml of basic fibroblast growth factor (bFGF; R&D systems). The colonies were passaged in clumps every 7 days using collagenase and mechanical dissociating methods.

### Differentiation of hES cells into otic neural progenitors (ONPs) and sensory neurons

Generation of otic neuroprogenitors was performed as described in Chen et al 2012. Briefly, undifferentiated hESCs were dissociated to single cells using 0.025% trypsin-EDTA (Sigma). After trypsinization, cell suspension was pelleted at 157 rcf for 5 min and passed through a 100 μm cell strainer (BD Labware) to get completely dissociated cells or very small aggregates. Cells were counted and plated onto laminin-coated T12.5 flasks (R&D systems, 5μg/cm^2^) at the seeding density of 4000 cells/cm^2^. Cells were maintained in a 1:1 mixture of Dulbecco’s modified Eagle’s medium (DMEM) and F12 nutrient supplemented with N2 and B27 (DFNB). FGF3 (50ng/ml) and FGF10 (50ng/ml) were added to the medium as supplements to drive the differentiation ^4,22^. The differentiated medium was replaced every 2 days. After 3-4 days, other epithelial-like and surrounding cells were scraped off to enrich for ONPs. The flask was cleaned from time to time and ONPs were then allowed to grow for further 10-12 days. The purified ONPs were then maintained in otic stem cell full medium or OSCFM (DFNB supplemented with bFGF (20ng/ml), IGF1 (50ng/ml) and EGF (20ng/ml)) and expanded on 0.1% gelatin-coated flask until sufficient cell numbers were obtained.

Neuralisation of ONPs into auditory neurons was induced with Shh-C24II (500ng/ml), bFGF (20 ngml) and neurotrophins (NT3 and BDNF; 10 ng/ml each) added at different time points as previously described by Chen et al. 2009 and 2012 ^4,22^.

### Preparation of polylactic acid electrospun scaffolds

Ten percent wt. of poly(L-lactide) (PLLA) were prepared by dissolving in 3:1 chloroform: methanol respectively. The solution was stirred for 1 hour at room temperature or until the polymer was totally dissolved. Three ml of polymer solution was electrospun from a 3 ml plastic syringe that attached to a flat tip needle at constant flow rate, which was controlled by a multispeed syringe pump. A high voltage (~10-12kv) was applied to the needle tip to charge the polymer solution and form a liquid jet. An aluminum-rotating disk was used as a grounded collector to collect the electrospun fibres. Random and high alignment of fibres was collected at the rotation speed of 180 and 2000 rpm respectively.

### Scanning electron microscopy (SEM)

The morphology and alignment of fibres and cells on the scaffolds were visualised by SEM (Philips XL-20) at an accelerating voltage of 20 kV. Each scaffold was coated with gold using a sputter coater (Emscope SC 500) prior to the observation. The images were then analysed using image J software (NIH).

### ONP cell culture on the scaffolds

The hES-derived ONP cells were seeded onto electrospun scaffolds that were sterilized in cold 70% ethanol for 15 min, washed 3 times with cold PBS and incubated in OSCFM medium overnight. About 8000 ONPs were seeded onto a 1 cm^2^ scaffold membrane, using a very small volume (20 μl). To allow attachment to the scaffold, cells were allowed to settle for one hour before the medium was topped up. ONPs were then maintained in either progenitor (OSCFM) or neuralising conditions for up to 14 days. Size of scaffolds and cell density were scaled up for qRT-PCR gene expression analysis (5×10^5^ cells/ 4 cm^2^ scaffold) in order to obtain good RNA yields.

### SEM sample preparation for ONPs on 3D scaffolds

As ONPs are loosely attached cells and most of them were not preserved under normal preparation, a new preparation protocol was established especially for ONPs. Culture medium was removed from the wells and the cells were fixed with warm Karnovsky’s fixative, which is a mixture of 2% paraformaldehyde, 2.5% glutaraldehyde and 0.1M Sodium Phosphate buffer, for 3 hours. ONPs were then dehydrated sequentially in 10%, 30% and 50% methanol for 5 min each, followed by an air dried for 3 days. The samples were dried again in a vacuum oven at room temperature overnight.

### Actin Staining

After the cells were fixed in 4% paraformaldehyde in PBS for 1 hr, they were rinsed with PBS and permeabilised with 0.3% Triton X-100 (PBST) for 15 min. Three hundred and fifty μl of Texas-red-X phalloidin (1:150, Invitrogen) was then added to each well and left on the rocking plate for 1 hr. The cells were then rinsed with PBS after phalloidin was removed and cell nuclei were counterstained with DAPI (1:200; BioLegend)

### Quantification of cell directionality

Fluorescent images from actin staining were used for the quantification of cell directionality. The images in red channel were converted to binary images using the thresholding method and the orientation of cells on binary images were then analysed by orientationJ distribution, an imageJ’s plugin for directional analysis. The distribution of orientation histogram was automatically generated from each image.

### Calcein-AM staining

Calcein-AM powder (Santa Cruz) was resuspended with anhydrous DMSO to make a 2mM stock solution. Working solution was prepared immediately prior to use by mixing Calcein-AM stock solution with the culture medium in 1:2000 dilutions. After the media was discarded, Calcein-AM working solution was added to each well and incubated under 5% CO_2_ at 37°C for 45 min. The cells were then washed with media and visualised under the fluorescent microscope. Intracellular esterases in live cells hydrolyse a non-fluorescent Calcein-AM to strongly fluorescent Calcein, which appeared green in cell cytoplasm. Numbers of Calcein positive cells from at least 10 different fields were used to calculate the estimated cell number on a 1cm^2^ scaffold.

### BrdU Viability assay

DNA synthesis monitoring immunofluorescence assay, 5-Bromo-2′ - deoxy-uridine (BrdU) labeling and detection kit I (Roche) was used to verify the viability and proliferative capacity of ONPs on the scaffolds. Ten μM BrdU was added to the culture medium and incubated for 48 hr before they were fixed with ethanol fixative, labelled with mouse anti-BrdU and anti-mouse fluorescein. The concentration of reagents and antibodies were used according to the manufacturer’s instructions. Viable cells were then quantified under fluorescent microscope.

### Immunolabelling and quantitation of fluorescence intensity

After 14 days on the scaffolds, the ONPs were fixed with 4% paraformaldehyde in PBS for 1 hr and permeabilised in 0.1%Triton X-100 for 15 min. Then, they were incubated for 1 hr at room temperature in blocking solution (0.1%Triton X-100, 5% normal donkey serum, 1% BSA in PBS) before the primary antibodies were added. Double staining of β-tubulin III (TUJ1) (1:100, mouse monoclonal, Covance) and NF200 (1:100, rabbit polyclonal, Sigma), as well as Tau (1:50, rabbit polyclonal, Santa Cruz) and MAP2 (1:200, mouse polyclonal, Abcam) were performed in this study. After the cells were incubated with primary antibodies overnight at 4°C, the antibodies were rinsed off and secondary donkey anti-mouse or anti-rabbit antibodies conjugated with Alexa Fluor 488 or Alexa Fluor 568 (1:250, Invitrogen) were applied and incubated for 1 hr at room temperature. For nuclei labeling, the cells were counterstained with DAPI (1:200, 4’,6- diamidino-2-phenylindole; BioLegend).

Fluorescence intensity per cell was evaluated using ImageJ software (NIH). All images were converted to black and white colour to avoid the bias. Freehand selection was used to draw the area over each cells and the level of fluorescence in a given region was measure as a mean grey value. This number was then subtracted by the average background intensity of that image.

### RNA extraction and gene expression analysis

Total RNA was extracted using Trizol (Invitrogen) and Qiagen RNeasy column combination protocol ^23^. 1 ml of Trizol was added to two 4 cm^2^ scaffolds and pipetted up and down to dissolve the fibres. After 5 min at room temperature, 200 μl of chloroform was added and mixed thoroughly. Each tube was incubated for 2 min at room temperature, then spin down at 12,000 g for 15 min at 4°C. After the centrifugation, the transparent upper phase was gently removed and transferred to a new tube. RNase-free 70% ethanol (500 μl was then added and mixed well using the pipette. The mixture was transferred to the RNeasy mini spin column and preceded according the manufacturer protocols.

QuantiTect Reverse Transcription Kit (Qiagen) was used to transcribe the extracted RNA to cDNA. Then, to check the expression of neural markers, real-time PCR was performed using Rotor-Gene SYBR Green (Qiagen). Primer sequences of housekeeping and target genes (from 5’ to 3’) were as follows: GAPDH, ATGGGGAAGGTGAAGGTCG and TAAAAGCAGCCCTGGTGACC; NTNG1, TTGTGGATTGGAAAGGCTGC and GGTTGGAAGGGATTCAGGGA; NPR2, AACGCCATGCACCAGAAATT and CATCTTCAGGCCAACAACCC; MMP13, TTGAGCTGGACTCATTGTCG and TCTCGGAGCCTCTCAGTCAT.

### Statistical analysis

All experiments were analysed using analysis of variance (one-way ANOVA) followed by Bonferroni’s post hoc comparison tests. Statistically significant differences will be considered if the p-value is less than 0.05 (p<0.05). All values are shown as mean ± standard deviation from at least 2 independent experiments. All the graphs and statistic calculations were performed on GraphPad Prism 6 (GraphPad Software, Inc., San Diego, CA, USA).

## RESULTS

### Visualization and quantification of fibrous scaffolds

After the fibres were fabricated, three independent electrospun scaffolds were visualized and quantified. SEM micrograph and histogram of angle difference demonstrated that most the fibres were aligned, with an average diameter of 0.93±0.1 μm (Figure. 1A and B). Random fibres were also fabricated and characterised to be used as a control in the experiments (Figure 1C). They were spun with the same set up as in aligned-fibres but the collector was rotated at the lowest speed. The distributions were almost equal in all angles for random fibres (Figure 1D).

**Figure 1.**
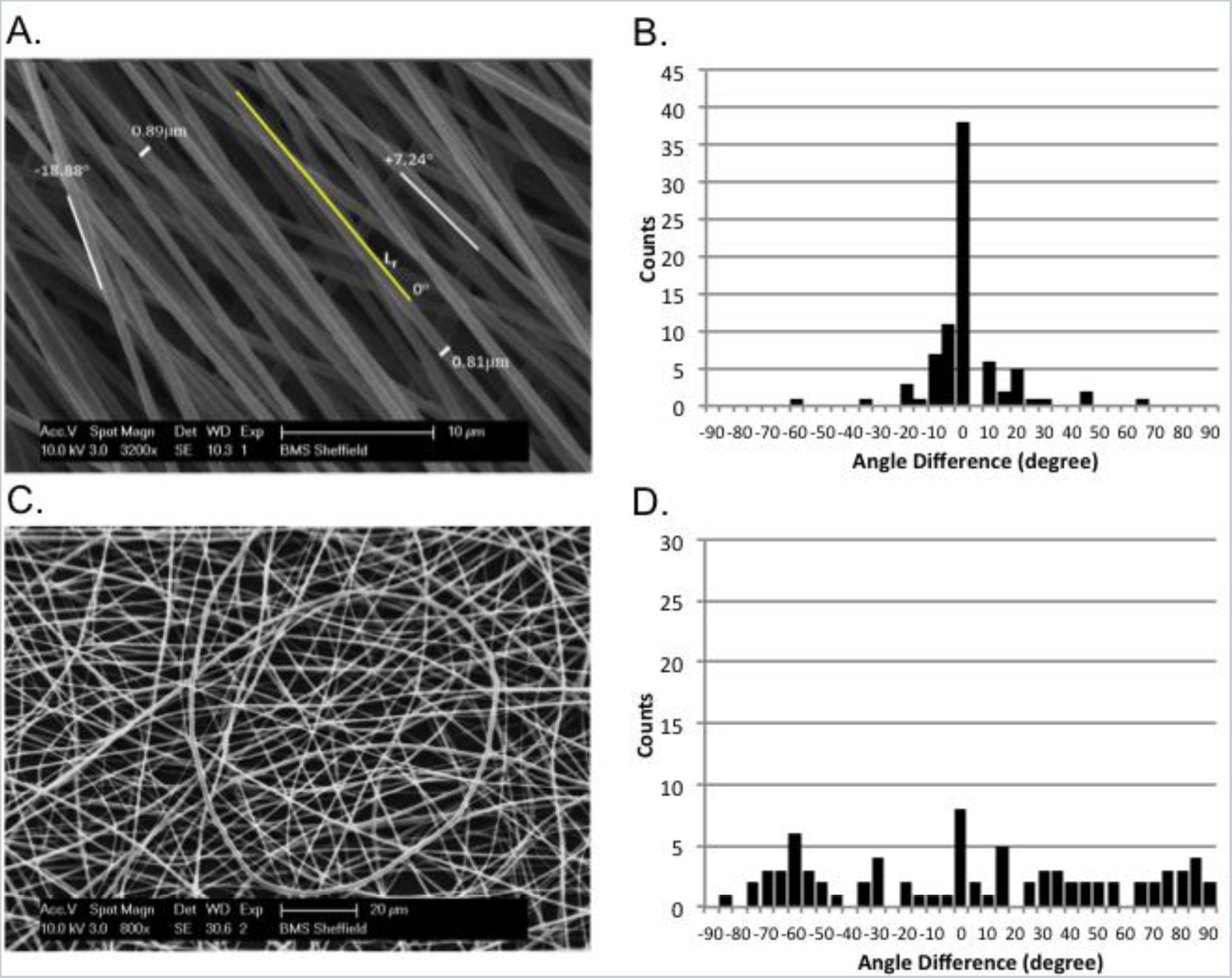
Electrospun fibres from optimized electrospinning parameters. (A) SEM image of PLLA aligned fibres and alignment analysis (scale bar = 10 μm). (B) Histogram of angle difference showing that most of the fibres are parallel to the reference line. (C) SEM image of random fibres collected with the lowest collector rotation speed (scale bar = 20 μm). (D) Histogram of angle difference on random fibres.

### The effect of aligned PLLA fibres on ONPs morphology

The morphologies of hESC-derived ONPs grown under progenitor conditions (OSCFM) for 3, 7 and 14 days on aligned PLLA fibres were investigated by actin staining (Texas-red Phalloidin). Nuclei were counterstained with DAPI. When cultured on tissue culture plastic for a few passages, ONPs were flat and spread with an average size of 300 μm (Figure 2A). On the contrary, a distinct morphology was observed on 3D fibres, with cells being smaller with an average size of 200 μm. After 3 days in culture, some of the ONPs were spreading on the fibres (arrowhead) and elongated cells were noticed (arrow) (Figure 2B). As for day 7 and 14, most of ONPs exhibited bipolar morphology (69±4%) and displayed a parallel, aligned pattern in some areas of the scaffold (arrow) (Figure 2C). However, cells with polygonal morphology were also presented (arrowhead). In some other areas on the scaffolds, randomness of ONP cell arrangement still existed (Figure 2D). On the other hand, ONP growing on the random PLLA fibres were spreading in all directions and showed irregular morphology (Figure 2E).

**Figure 2.**
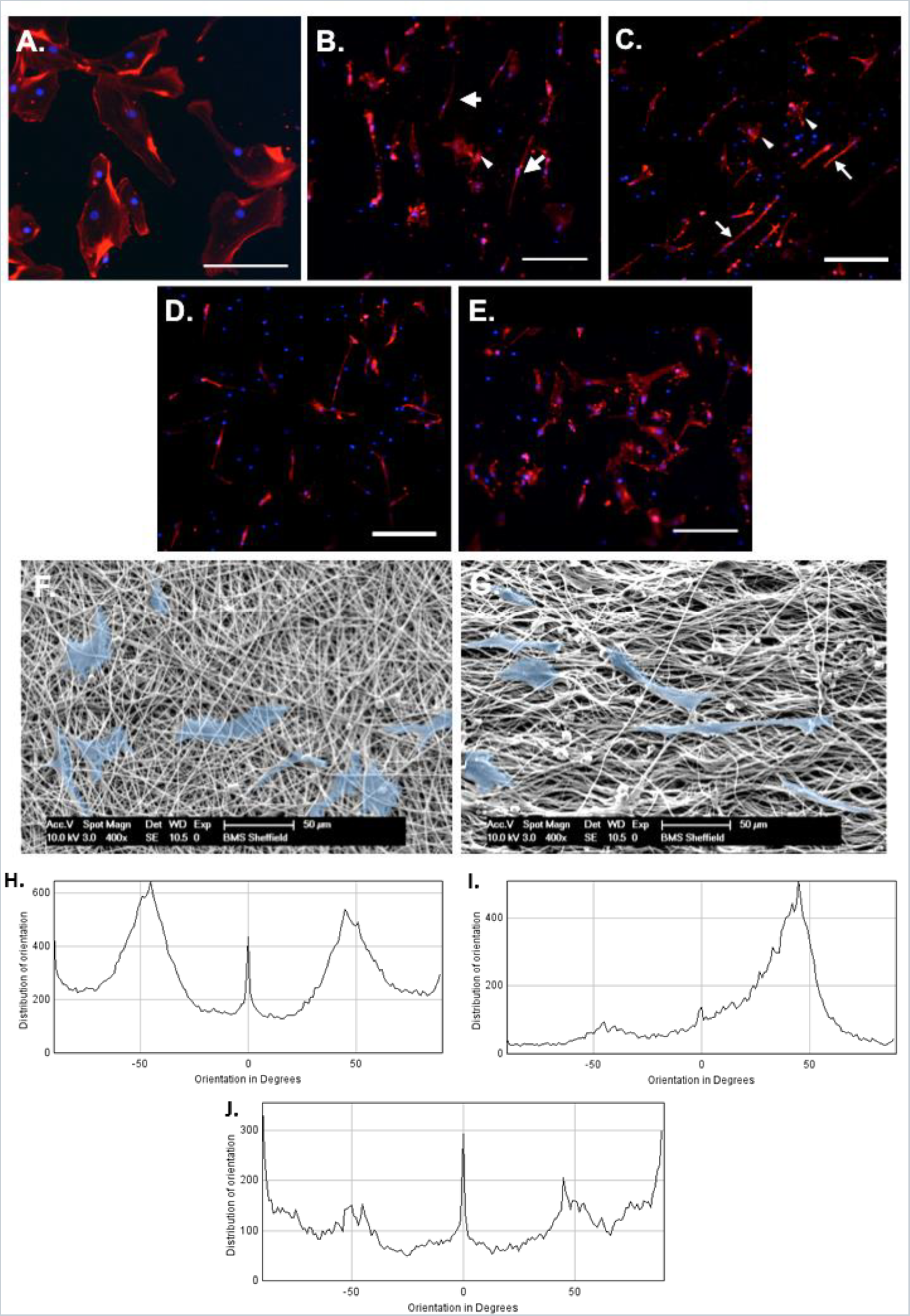
Actin staining showing ONP cell morphology on **(A)** tissue culture plastic **(B)** PLLA aligned fibres after 3 days in culture **(C,D)** PLLA aligned fibres after 14 days in culture **(E)** PLLA random fibres after 14 days in culture. Scale bar=200μm. Scanning electron micrographs of the cells were shown on **(F)** random and **(G)** aligned PLLA fibres. The cells were masked in blue for the visibility. Scale bar=50μm. Distribution of cell orientation on tissue culture plastic, PLLA aligned and random fibres were measured and shown as histograms on (H), (I) and (J) respectively.

SEM is another method that was used to confirm the findings from fluorescent imaging. ONPs indeed extend along the fibre axes and the cells with different orientation or irregular shape arise from the crossing fibres presence in the scaffolds (Figure 2F,G). The micrographs also revealed that single ONP cells attached to few fibres at a time.

Orientations of cell distributions were quantified using OrientationJ plugin from ImageJ. The orientation of each pixel is evaluated and plotted as a histogram against the orientations in degree. The histograms demonstrated that ONPs on 2D tissue culture plastic (Figure 2H) and PLLA random fibres (Figure 2J) have broad distributions of orientation and several peaks were presented in the histograms. While the cells on PLLA aligned fibres (Figure 2I) exhibited one big peak, which indicated that most of the cells were arranged in a certain degree.

### The effect of aligned PLLA fibres on ONPs viability and proliferation

To check the number of viable ONP cells on the scaffolds, Calcein-AM staining was carried out. After 14 days of culture, the number of viable cells on neuralising 2D, control (OSCFM) PLLA and neuralising PLLA were all significantly lower than those found on control (OSCFM) 2D (Figure 3A). The lower numbers found on PLLA could be partially explained by the lower cell attachment expected on PLLA fibres, as no bioactive sites are presented and they were not treated with any chemicals or biological compounds that would enhance cell attachment. A decreased cell number under neuralising condition could also infers that ONPs are differentiating into non-proliferating neurons ^24^. Therefore, the population of cells with dividing capability is lower. It is noted that the differences between control and neuralising conditions on 2D were larger than those on PLLA.

**Figure 3.**
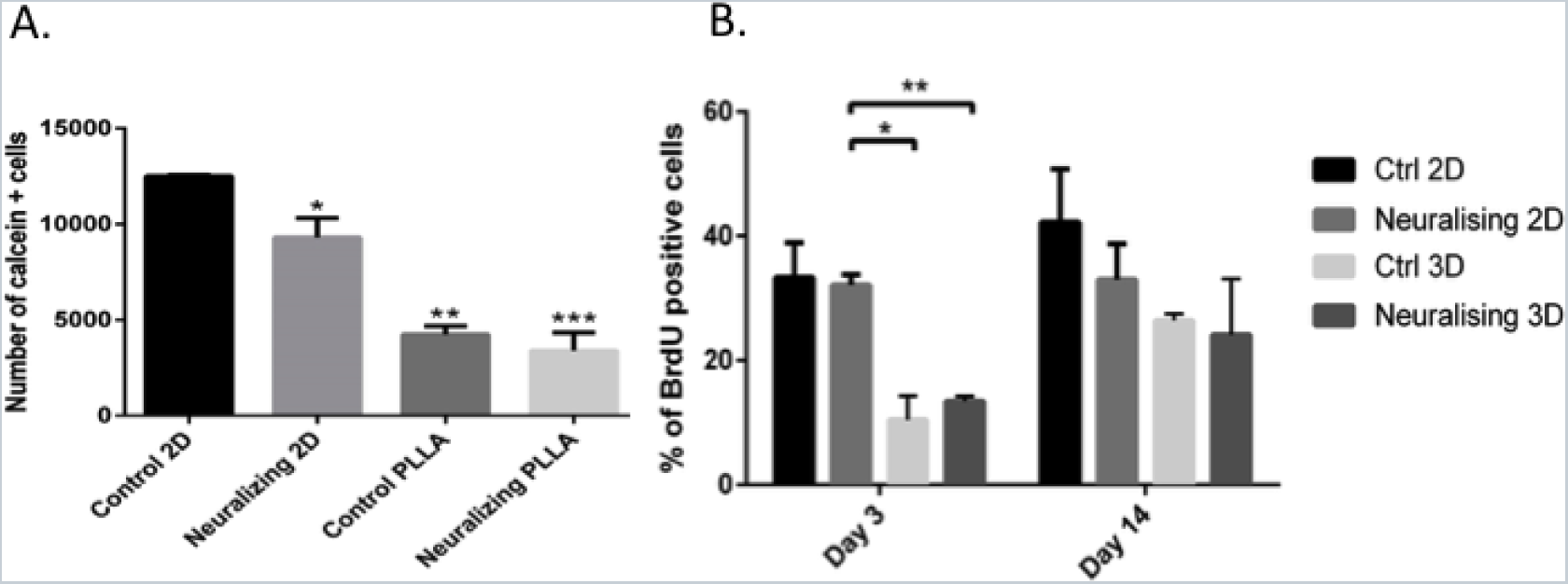
Viability and proliferation of ONPs on PLLA fibres. (A) Number of viable ONPs on 1×1cm^2^ PLLA aligned fibres after 14 days in culture. Data is mean±SD. (B) ONPs proliferation after 3 and 14 days culture on PLLA aligned fibres. *p<0.05, **p<0.01

The proliferation of ONPs was also investigated using a BrdU incorporation assay (figure 3B). A significant reduction of cell proliferation, triggered by culturing cells on PLLA was noticed after 3 days, regardless of the culture conditions. The trend was maintained after 14 days, although the differences were not large enough to remain significant. The reduction of the percentage of BrdU positive cells in neuralising conditions was, however, more obvious.

### The effect of aligned PLLA fibres on ONPs differentiation

We investigated the differentiation behaviour of ONPs on PLLA electrospun fibres in both conditions (progenitor and neuralising) using immunostaining of the early neuronal marker β-tubulin III (TUJ1), and the mature neural marker neurofilament-200 (NF200). After 14 days in culture, the expressions of TUJ1 and NF200 were enhanced when the cells were grown in the neuralising condition or on aligned fibres (noted that cell number per area on the fibres are lower than those on 2D) (Figure 4A). Mean Fluorescent intensity per cell and number of positive cells for both markers were increased even without any neurotrophins supplement (Figure 4B). On 2D, the intensities of both markers were relatively the same in control and neuralising conditions but numbers of positive cells were higher in the neuralising one. Similar trend was observed on random fibres with a slight increase of fluorescence intensity. The significant difference was detected only on TUJ1 between control 2D and neuralising random fibres.

**Figure 4.**
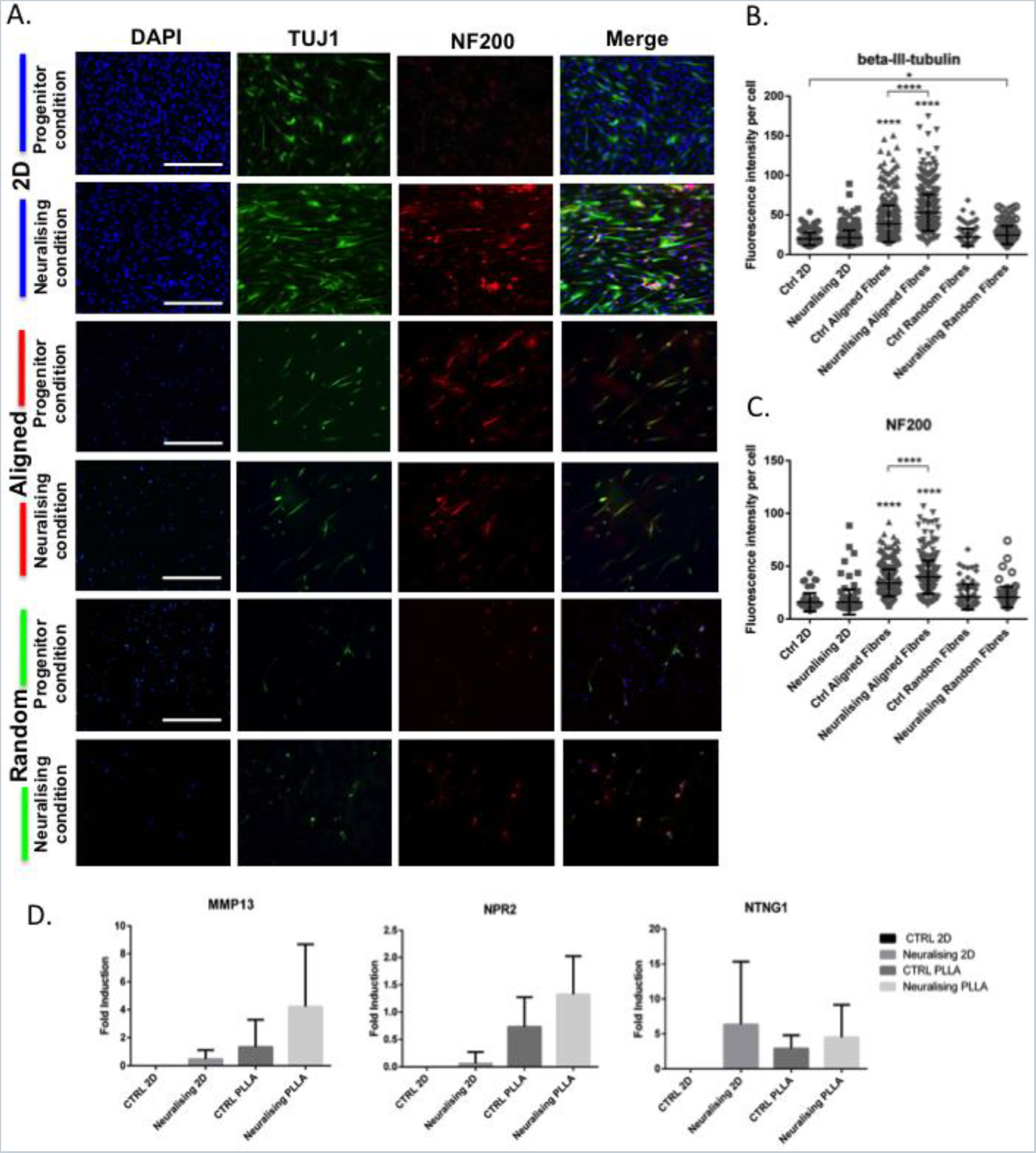
The differentiation of ONPs on PLLA fibres. (A) Immunostaining of ONPs on 2D tissue culture plastic and 3D aligned PLLA fibres for neuronal markers, TUJ1 or ß-tubulinIII (green) and NF200 (red). Scale bar=400 μm. (B) Mean fluorescent intensity per cell of ß-tubulinIII and (C) NF200 on different culture materials. *p<0.05, ****p<0.0001 (D) The effect of aligned substrates on the expression of MMP13, NPR2 and NTNG1 in ONPs. Data is mean±SD.

Expression of both neuronal markers studied was enhanced on ONPs seeded on aligned fibres under both culture conditions. There was a clear, statistically significant shift of labelling intensities (p<0.0001). This indicated that the alignment of fibres alone could effectively drive the differentiation of ONPs. Neuralising condition increased the neuronal marker expression on aligned fibres even further and the intensity of both TUJ1 and NF200 were significantly increased comparing to control condition.

Several molecular markers known to be highly expressed in differentiated ONPs were also explored. Matrix metalloproteinase (MMP13), natriuretic peptide receptor2 (NPR2) and Netrin-G1 (NTNG1) ^4^, were used as differentiation marker genes and analysed by qRT-PCR (Figure 4C). These genes are upregulated in a specific manner during SGN development. MMP13, also known as collagenase-3, is a zinc-dependent proteolytic enzyme involved in the remodeling of ECM and is enriched at late developmental stage of SGN (Lu et al., 2011; NCBI, 2016a). *NPR2* encodes an integral membrane receptor for natriuretic peptides, which is required for axon bifurcation in SGN and auditory circuit assembly ^25,27,28^. NPR2 is highly expressed at very early stage, then dropped in the middle and finally raised again in the late developmental stage of SGN. *NTNG1* encodes a preproprotein that can be further processed into a secreted protein, which is responsible for axon growth and guidance. It is highly expressed throughout the SGN development but the level is slightly higher in the early stage ^25,29^.

All three genes were upregulated when ONPs were cultured in neuralising medium and on the aligned scaffolds. The increment ranged from 1 to 6 folds. The upreguration of MMP13 and NPR2 on aligned fibres were even higher than those on 2D neuralising conditions. On the contrary, the induction of NTNG1 was maximised on 2D neuralising condition. However, due to wide deviations, none of them are considered as significantly different. From the pattern of gene expression that were observed, it can be inferred that after 14 days of differentiation, ONPs on aligned fibres are in a late developmental stage.

### The effect of aligned PLLA fibres on ONPs polarization

To verify the polarisation of differentiated cells, ONPs were fixed and co-stained with anti-TAU (axonal marker) and anti-MAP2 (dendritic marker) antibodies after 14 days under different culture conditions. On TC plastic, ONPs under progenitor condition rarely expressed TAU and MAP2 but a slight increase of TAU expression was observed and cells with positive MAP2 were identified under neuralising condition (Figure. 5A). On align fibres, ONPs in both progenitor and neuralising conditions were positive for both neuronal polarity markers. Some ONPs also showed specific localization of MAP2 in the soma and TAU in both soma and axon indicating a polarization of those cells (arrow). However, the majority of the cell population did not yet specify their neuronal polarity. The number of positive cells was then quantified and the results revealed that neuralising 2D, control PLLA and neuralising PLLA can increase the percentage of TAU and MAP2 positive cells by 2 to 8 folds when comparing to Control 2D group (Figure 5B). The percentages of those 3 conditions were comparable but the most prominent one was observed on neuralising PLLA.

**Figure 5.**
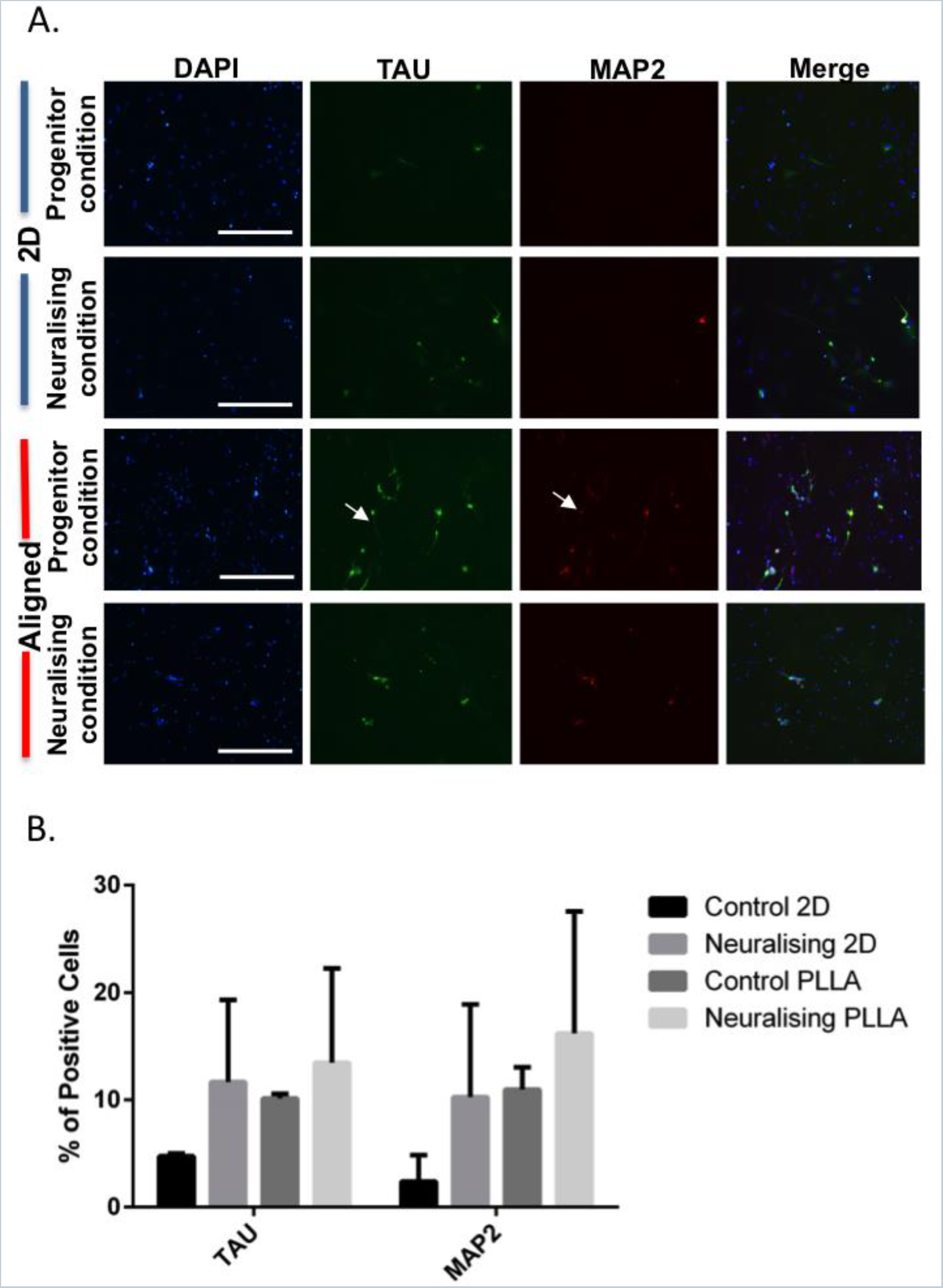
ONP polarization on PLLA aligned fibres. (A) Immunolabelling of neural polarity markers, TAU (green) and MAP2 (red) on ONPs. Scale bar=400 μm. (B) Percentage of TAU and MAP2 positive ONPs when cultured on different substrates for 14 days. Data shown are mean±SD.

## Discussion

Not only limited to mature cells, the correlation between substrate properties or microenvironment and stem cell fate has been widely investigated. Stem cell differentiation is one of the main interests in tissue engineering and regeneration, as in many cases the desired cell type is required prior to transplantation for the safety and efficiency of the treatment ^30^. It is increasingly clear that the native microenvironment that supports stem cells or ‘stem cell niche’ plays a crucial role in the control of stem cell behaviour and fate decision ^31,32^. Thus, a considerable amount of work is currently being performed employing this view that allows taking control over stem cells and directing them to the desired behaviour.

Numerous studies have explored the use of electrospun fibres as tissue engineering scaffolds for the repair of both central and peripheral nervous system injuries ^33,34^. Aligned electrospun fibres have also been used by many research groups. Its advantages in neurite guiding and extension has been reported *in vitro* ^35–37^ and the success repair were also demonstrated *in vivo* ^38,39^. However, those studies were performed with the cells that are not directly relevant to the auditory system and tissue engineering for SGN damage or degeneration has never been tried. Here, we investigated the response of ONPs, a promising cell type that could be used for SGN regeneration, to the aligned fibrous tissue-engineered substrate (PLLA).

There are many parameters that affect electrospinning process and fibre morphology. They are needed to be optimised in order to achieve the ideal scaffolds for each work ^40^. So, in order to obtain an aligned fibre mesh with the diameter of ~1 μm, which is expected to be the compromise fibre size for neuronal guidance, proliferation and differentiation ^41,42^, electrospinning parameters were manipulated. The most suitable parameters obtained were used to produce aligned and random electrospun fibres. The visualisation and quantification of SEM images showed that we can successfully obtain both fibre types.

One of the very first ONP features that we monitored when grown on PLLA fibres was cell morphology. The dissimilarity of ONPs morphology on 2D and 3D matrices are to be expected as the cells that were cultured on 2D tissue culture plastic are flat and can only spread in horizontal plane, while on 3D, which is considered to be closer to physiological conditions, cells can spread in all directions ^43^. The parallel cell pattern is a promising result indicating that alignment of the fibres can act as a guidance cues for ONPs. Calcein-AM staining and BrdU assay also demonstrated that the numbers of viable, attached cells and the percentage of cell proliferation were lower on the fibres comparing to 2D culture plastic. These findings are coherent with the idea cells are differentiating, reducing or stopping proliferation altogether ^24^.

In previous studies of the role of scaffolds on neural differentiation, experiments have been performed under neural-inducing conditions such the presence of neurotrophins. However, the effect of the external scaffold on the progenitors is rarely described ^44–46^. From the expression of neuronal markers in protein and RNA level, we clearly showed that ONPs were sensitive to the change of physical environment (aligned fibres) and were able to differentiate even in the normal expansion medium. Under neuralising condition, ONP differentiation was increased even further. This indicated that there is a synergistic effect between physical (fibre alignment) and chemical cues (neuralising agents).

In order for neural cells to function after they are differentiated, polarization and/or establishment of axons and dendrites are required ^47^. Beside from intrinsic factors, extracellular signal also controls polarity specification as the neural cells are physiologically surrounded by extracellular matrix. Various axonal guidance and repellent molecules have been identified to effectively control the asymmetric growth of neurites and leads to axon or dendrite formation ^48,49^. Some growth factors such as NT3 and BDNF that were used in our differentiation medium also help enhancing neuronal polarization ^50,51^. As we previously found that ONPs are differentiated on the aligned fibres and lots of them are positive for mature neuronal marker (NF200), the establishment of neural polarity is another aspect to confirm the transition to a mature neuronal phenotype. Culturing of ONPs on aligned scaffolds and under neuralising condition increased the expression of neuronal polarity markers (TAU and MAP2) but the polarities of those positive cells were not yet to be specified.

## CONCLUSIONS AND FUTURE WORKS

In summary, we have demonstrated that otic neural progenitors (ONPs) can attach, survive and be able to differentiate on aligned PLLA fibres. The physical cue (alignment of fibres) can help enhancing the differentiation of ONPs even in the absence of neuralising agents and guiding neurite extension along the alignment axis. In addition, it also influences changes on ONP morphology and the specification of neural polarity. This aligned PLLA scaffolds can also be further manipulated to facilitate the transplantation. A conduit could be made by rolling the PLLA electrospun sheet and seed it with ONPs and glial cells. These applications could be highly useful to develop new strategies to repair the auditory nerve.

## ACKNOWLEDGEMENT

The authors would like to thank Dr. Priya Vishwanathan and Rivolta’s lab members for their kind help in teaching new lab techniques. This work was supported by the Royal Thai Government and partially supported by MRC award to Prof. Marcelo N. Rivolta.

## CONTRIBUTIONS

All authors participate in the designed and interpretation of the studies. KN performed the experiments, analysed the data and wrote the manuscript. MNR and GB contributed to the revision of the manuscript.

## COMPETETING INTERESTS

The authors declare no conflict of interest.

## REFERENCES

1. Groves, A. K., Zhang, K. D. & Fekete, D. M. The genetics of hair cell development and regeneration. Annu. Rev. Neurosci. 36, 361–81 (2013).

2. Hu, Z. et al. Survival and neural differentiation of adult neural stem cells transplanted into the mature inner ear. Exp. Cell Res. 302, 40–47 (2005).

3. Incesulu, A. & Nadol, J. Correlation of acoustic threshold measures and spiral ganglion cell survival in severe to profound sensorineural hearing loss: Implications for cochlear implantation. Ann. Otol. Rhinol. Laryngol. 107, 906–911 (1998).

4. Chen, W. et al. Restoration of auditory evoked responses by human ES-cell-derived otic progenitors. Nature 490, 278–282 (2012).

5. Lang, H., Schulte, B. A. & Schmiedt, R. A. Ouabain induces apoptotic cell death in type I spiral ganglion neurons, but not type II neurons. JARO - J. Assoc. Res. Otolaryngol. 6, 63–74 (2005).

6. Burdick, J. A. & Mauck, R. L. Biomaterials for Tissue Engineering Applications. (Springer-Verlag Wien, 2011).

7. Discher, D. E., Mooney, D. J. & Zandstra, P. W. Growth factors, matrices, and forces combine and control stem cells. Science 324, 1673–7 (2009).

8. Shibata, S. B. & Raphael, Y. Future approaches for inner ear protection and repair. Journal of Communication Disorders 43, 295–310 (2010).

9. Clarke, J. C. et al. Micropatterned methacrylate polymers direct spiral ganglion neurite and Schwann cell growth. Hear. Res. 278, 96–105 (2011).

10. Mattotti, M., Micholt, L., Braeken, D. & Kovačić, D. Characterization of spiral ganglion neurons cultured on silicon micro-pillar substrates for new auditory neuro-electronic interfaces. J. Neural Eng. 12, 26001 (2015).

11. Reich, U. et al. Differential fine-tuning of cochlear implant material-cell interactions by femtosecond laser microstructuring. J. Biomed. Mater. Res. - Part B Appl. Biomater. 87, 146–153 (2008).

12. Stöver, T. & Lenarz, T. [Biomaterials in cochlear implants]. Laryngorhinootologie. 88 Suppl 1, S12–S31 (2009).

13. Evans, A. R. et al. Laminin and fibronectin modulate inner ear spiral ganglion neurite outgrowth in an in vitro alternate choice assay. Dev. Neurobiol. 67, 1721–1730 (2007).

14. Kikkawa, Y. S. et al. Growth factor-eluting cochlear implant electrode: impact on residual auditory function, insertional trauma, and fibrosis. J. Transl. Med. 12, 280 (2014).

15. Paasche, G. et al. Technical report: modification of a cochlear implant electrode for drug delivery to the inner ear. Otol. Neurotol. 24, 222–7 (2003).

16. Pettingill, L. N., Richardson, R. T., Wise, A. K., O’Leary, S. J. & Shepherd, R. K. Neurotrophic factors and neural prostheses: Potential clinical applications based upon findings in the auditory system. IEEE Transactions on Biomedical Engineering 54, 1138–1148 (2007).

17. Ryan, A. F., Wittig, J., Evans, A., Dazert, S. & Mullen, L. Environmental micropatterning for the study of spiral ganglion neurite guidance. Audiol. Neurotol. 11, 134–143 (2005).

18. Stover, T. et al. Development of a drug delivery device: using the femtosecond laser to modify cochlear implant electrodes. Cochlear Implant. Int 8, 38–52 (2007).

19. Xie, J. et al. Neurotrophins differentially stimulate the growth of cochlear neurites on collagen surfaces and in gels. NEURAL Regen. Res. 8, (2013).

20. Schwieger, J. et al. Neuronal survival, morphology and outgrowth of spiral ganglion neurons using a defined growth factor combination. PLoS One 10, (2015).

21. Santi, P. A. & Johnson, S. B. Decellularized ear tissues as scaffolds for stem cell differentiation. JARO - J. Assoc. Res. Otolaryngol. 14, 3–15 (2013).

22. Chen, W. et al. Human fetal auditory stem cells can be expanded in vitro and differentiate into functional auditory neurons and hair cell-like cells. Stem Cells 27, 1196–1204 (2009).

23. Untergasser, A. RNAprep - Trizol combined with Columns. Untergasser’s Lab (2008). at <http://www.untergasser.de/lab/protocols/rna_prep_comb_trizol_v1_0.htm>

24. Purves, D. et al. Neuroscience. (Sinauer Associate, Inc., 2008). doi:978-0878937257

25. Lu, C. C., Appler, J. M., Houseman, E. A. & Goodrich, L. V. Developmental Profiling of Spiral Ganglion Neurons Reveals Insights into Auditory Circuit Assembly. J. Neurosci. 31, 10903–10918 (2011).

26. NCBI. MMP13 matrix metallopeptidase 13 [Homo sapiens (human)]. (2016). at <http://www.ncbi.nlm.nih.gov/gene/4322>

27. Lu, C. C. et al. Mutation of Npr2 Leads to Blurred Tonotopic Organization of Central Auditory Circuits in Mice. PLoS Genet. 10, (2014).

28. NCBI. NPR2 natriuretic peptide receptor 2 [Homo sapiens (human)]. (2016). at <http://www.ncbi.nlm.nih.gov/gene/4882>

29. NCBI. NTNG1 netrin G1 [Homo sapiens (human)]. (2016). at <http://www.ncbi.nlm.nih.gov/gene/22854>

30. Guillot, P. V, Cui, W., Fisk, N. M. & Polak, D. J. Stem cell differentiation and expansion for clinical applications of tissue engineering. J. Cell. Mol. Med. 11, 935–44 (2007).

31. Schofield, R. The relationship between the spleen colony-forming cell and the haemopoietic stem cell. Blood Cells 4, 7–25 (1978).

32. Li, L. & Xie, T. Stem cell niche: structure and function. Annu. Rev. Cell Dev. Biol. 21, 605–631 (2005).

33. Liu, W., Thomopoulos, S. & Xia, Y. Electrospun nanofibers for regenerative medicine. Adv. Healthc. Mater. 1, 10–25 (2012).

34. Xie, J., MacEwan, M. R., Schwartz, A. G. & Xia, Y. Electrospun nanofibers for neural tissue engineering. Nanoscale 2, 35–44 (2010).

35. Schnell, E. et al. Guidance of glial cell migration and axonal growth on electrospun nanofibers of poly-εcaprolactone and a collagen/poly-[epsilon]-caprolactone blend. Biomaterials 28, 3012–3025 (2007).

36. Shen, Y. et al. Guidance of olfactory ensheathing cell growth and migration on electrospun silk fibroin scaffolds. Cell Transplant. 19, 147–157 (2010).

37. Xie, J. et al. The differentiation of embryonic stem cells seeded on electrospun nanofibers into neural lineages. Biomaterials 30, 354–362 (2009).

38. Hurtado, A. et al. Robust CNS regeneration after complete spinal cord transection using aligned poly-l-lactic acid microfibers. Biomaterials 32, 6068–6079 (2011).

39. Kim, Y. tae, Haftel, V. K., Kumar, S. & Bellamkonda, R. V. The role of aligned polymer fiber-based constructs in the bridging of long peripheral nerve gaps. Biomaterials 29, 3117–3127 (2008).

40. Bhardwaj, N. & Kundu, S. C. Electrospinning: A fascinating fiber fabrication technique. Biotechnology Advances 28, 325–347 (2010).

41. Binder, C., Milleret, V., Hall, H., Eberli, D. & Lühmann, T. Influence of micro and submicro poly(lactic-glycolic acid) fibers on sensory neural cell locomotion and neurite growth. J. Biomed. Mater. Res. - Part B Appl. Biomater. 101, 1200–1208 (2013).

42. Christopherson, G. T., Song, H. & Mao, H. Q. The influence of fiber diameter of electrospun substrates on neural stem cell differentiation and proliferation. Biomaterials 30, 556–564 (2009).

43. Baker, B. M. & Chen, C. S. Deconstructing the third dimension - how 3D culture microenvironments alter cellular cues. J. Cell Sci. 125, 3015–3024 (2012).

44. Gerardo-Nava, J. et al. Human neural cell interactions with orientated electrospun nanofibers in vitro. Nanomedicine (Lond). 4, 11–30 (2009).

45. Lim, S. H., Liu, X. Y., Song, H., Yarema, K. J. & Mao, H. Q. The effect of nanofiber-guided cell alignment on the preferential differentiation of neural stem cells. Biomaterials 31, 9031–9039 (2010).

46. Liu, X. et al. Guidance of neurite outgrowth on aligned electrospun polypyrrole/poly(styrene-beta-isobutylene-beta-styrene) fiber platforms. J. Biomed. Mater. Res. - Part A 94, 1004–1011 (2010).

47. Arimura, N. & Kaibuchi, K. Neuronal polarity: from extracellular signals to intracellular mechanisms. Nat. Rev. Neurosci. 8, 194–205 (2007).

48. Adler, C. E., Fetter, R. D. & Bargmann, C. I. UNC-6/Netrin induces neuronal asymmetry and defines the site of axon formation. Nat. Neurosci. 9, 511–518 (2006).

49. Whitford, K. L., Dijkhuizen, P., Polleux, F. & Ghosh, A. Molecular control of cortical dendrite development. Annu. Rev. Neurosci. 25, 127–49 (2002).

50. Da Silva, J. S., Hasegawa, T., Miyagi, T., Dotti, C. G. & Abad-Rodriguez, J. Asymmetric membrane ganglioside sialidase activity specifies axonal fate. Nat Neurosci 8, 606–615 (2005).

51. Huang, E. J. & Reichardt, L. F. Trk Receptors: Roles in Neuronal Signal Transduction. Annu. Rev. Biochem. 72, 609–642 (2003).

